# Socioeconomic variation across multiple cities predicts avian life-history strategies

**DOI:** 10.1101/2020.04.23.058537

**Authors:** Riikka P Kinnunen, Kevin Fraser, Chloé Schmidt, Colin J Garroway

**Affiliations:** Department of Biological Sciences, Biological Sciences Building, University of Manitoba, Winnipeg, MB, Canada R3T 2N2

**Author notes:** Correspondence: Riikka Kinnunen, Biological Sciences Building, University of Manitoba, 50 Sifton Rd, Winnipeg, MB, Canada R3T 2N2, Colin Garroway, Biological Sciences Building, University of Manitoba, 50 Sifton Rd, Winnipeg, MB, Canada R3T 2N2. Statement of authorship: RPK, CS, and CJG conceived of the study. RPK and KF collected data and RPK built the dataset. RPK performed statistical analyses with input from CS and CJG. All authors contributed to data interpretation. RPK and CJG wrote the first draft of the manuscript, and all authors contributed to subsequent revisions.

## Abstract

Cities are the planet’s newest ecosystem and thus provide the opportunity to study community formation directly following major permanent environmental change. The human social and built components of environments can vary widely across different cities, yet it is largely unknown how these features of a city covary with the traits of colonizing species. We constructed a new dataset from open-source data with 44,670 observations of 160 Passerine species observed 1,908 urban areas across the United States. We found that as a city’s housing density and median income increased it tended to support more migratory species and species with smaller body sizes and shorter lifespans. This suggests that differential survival and reproduction at the species-level can quickly generate geographical patterns of species trait variation across urban environments similar to those found in natural settings without the need for evolutionary change.

## Introduction

Spatial gradients of species life-history traits can occur if species with similar phenotypes tend to co-occur in similar environments (e.g., Vellend 2016; Pontarp et al. 2019). This is ecological selection (in the sense of Vellend 2016)—the phenotype-based differential survival and reproduction of members of different species due to environmental filtering (Kraft et al. 2015) and biotic interactions. Together with ecological drift, dispersal, and speciation, ecological selection contributes to the formation of spatial gradients in species richness, abundance, and composition (Vellend 2016). When environments change rapidly, ecological selection should be fast-acting with detectable effects in contemporary timeframes—species with sufficient access to a new environment either can or cannot persist in it. Ecological selection should be particularly important for shaping communities immediately following disturbances and setting the stage for future evolutionary change in the new environment. Here we use the creation of cities as an instance of new habitat formation suited to exploring contemporary ecological selection.

Cities are the planet’s newest ecosystems—ones where human social and built environments interact with natural physical and biological environments to shape ecosystem dynamics (Hobbs et al. 2006; Pickett et al. 2017). Cities cover ∼3% of habitable land globally (Liu et al. 2014) with continued growth forecasted (Seto et al. 2011). Thus, city building has created a natural experiment with which we can explore how ecological selection shapes biodiversity following the rapid creation of a major new habitat type. While we know that cities are causing evolutionary change in urban species (e.g., Johnson & Munshi-South 2017; Schmidt et al. 2020), the ecological processes that shape initial biodiversity patterns—the starting material for subsequent evolutionary change—are less well known. To explore this, we analyzed how trait variation in native Passerines breeding in cities in the United States varies with socioeconomic features of different cities. We focused on socioeconomic heterogeneity because it reflects different aspects of resource availability and environmental disturbance levels, two factors known to correlate well with life-history trait variation. A city’s socioeconomics directly reflects human activities—the defining feature of urban environments (Pickett et al. 2017). Despite this likely importance, the influence of socioeconomic heterogeneity on community formation is not well understood (Pickett et al. 2017).

Some species-level traits show clinal variation—thought to reflect adaptations to environmental variation (Lack 1947; James 1970; Evans et al. 2006; Olson et al. 2009). Resource availability and environmental stability in natural settings both tend to be negatively correlated with species body size and lifespan and positively related to reproductive output (Lack 1947, 1954; Martin 1987; Ricklefs 2000). This is thought to be because 1) large-bodied species are better able to withstand periods of low resource availability; 2) investment in fewer better-quality offspring in poor environments increases their likelihood of survival; 3) and being long-lived allows for multiple reproductive attempts given higher offspring mortality rates (Sol et al. 2012). Migration is another important life-history strategy associated with resource availability that allows species to cope with seasonality and periods of low productivity, particularly among birds, and migratory species generally select resource-rich stable habitats during the breeding season (Somveille et al. 2015).

There is now considerable evidence from both birds and mammals suggesting that cities filter for subsets of local species that have traits suited to population persistence in urban environments (Chace & Walsh 2006; Kark et al. 2007; Croci et al. 2008; Leveau 2013; Meffert & Dziock 2013; Meillère et al. 2015; Aronson et al. 2016; Silva et al. 2016; Jokimäki et al. 2016; Alberti et al. 2017; Leveau et al. 2017; Sepp et al. 2018; Santini et al. 2019; Hensley et al. 2019). Relative to non-urban species, successful urban colonizers tend to have larger body masses, bigger brains and be tolerant of a broad range of environments (Bonier et al. 2007; Croci et al. 2008; Maklakova et al. 2011; Lowry et al. 2013; Iglesias-Carrasco et al. 2019). The use of cities by migratory species is relatively unexplored, but there is some evidence that migratory birds may be underrepresented in urban ecosystems (Allen & O’Connor 2000; Kluza et al. 2000; Poague et al. 2000). Additionally, habitat suitability in cities represents a strong filter against tree and ground nesting birds compared to species that nest in artificial cavities, or on buildings (Lim & Sodhi 2004; Conole & Kirkpatrick 2011; Lizée et al. 2011; Jokimäki et al. 2016). Much of this previous work has focused on urban–rural species trait comparisons that treat different cities as homogeneous with respect to the features of species that can colonize them. However, it is notable that in instances where species filtering has been explored across multiple cities different cities seem to filter for slightly different subsets of traits (Hensley et al. 2019). This finding suggests that different cities may be suited to different life-history strategies.

While cities are more similar to each other than they are to nearby rural lands, there is also clear variation across different cities in both their built and human social components. For example, plant diversity and vegetation cover—important predictors of urban biodiversity—are higher in wealthier areas (Talarchek 1990; Iverson & Cook 2000; Hope et al. 2003; Kinzig et al. 2005; Leong et al. 2018). At more local scales avian biodiversity is affected by features such as urban canopy cover, the composition of landscape plantings and the presence of lawns, the prevalence of bird feeders, and human food waste (Thompson et al. 2003; Lepczyk et al. 2004; Smith et al. 2005, 2006; Tryjanowski et al. 2015). These features will be correlated with aspects of city-wide socioeconomic status and therefore biodiversity patterns across different urban areas. If ecological selection is important for community formation in urban environments we should expect that species best able to tolerate conditions in a particular city would share the phenotypic traits that enable persistence, just as we see trait variation converge across natural environments (Cleary et al. 2007; Jetz et al. 2008).

Here we ask whether socioeconomic heterogeneity across cities is predictive of the traits of Passerine species that colonize them. We used housing density, human population size, and median income as socioeconomic metrics likely to capture different axes of resource availability across cities. We hypothesized that wealthier cities would provide stable food and nesting resources similar to more natural and semi-natural resources in less disturbed contexts. We additionally hypothesized that cities with high housing densities and large human populations would provide more anthropogenic food resources due to supplemental feeding and food waste and might provide more nesting opportunities for species that nest on built structures or artificial cavities. However, highly populated, densely housed cities will also be the most disturbed, polluted, and fragmented—all factors that could modify the relationship between traits and resources. We consequently predicted that cities with higher median incomes would impose ecological selection pressures similar to stable, resource-rich environments, favoring small, short-lived species with high reproductive outputs. In contrast cities with high housing densities and large human populations, which are more disturbed and stochastic environments, would select for larger, long-lived species that invest more in offspring quality over quantity.

## Methods

### Data compilation

We compiled a new dataset from open data sources (see SI Fig 1 for detailed data compilation process). First, we downloaded the eBird Basic Dataset for the United States from eBird.org (Sullivan et al. 2009). eBird is an online bird abundance and distribution checklist program jointly coordinated by the Cornell Laboratory of Ornithology and the National Audubon Society. The eBird project relies on citizen science volunteer observers who submit georeferenced observations of species to a centralized database. Regional reviewers identify outliers and verify each species observation based on sighting coordinates (Wood et al. 2011). The eBird Basic Dataset is a publicly available large core dataset with >100,000,000 observations worldwide (available from https://ebird.org/). We used eBird observations of Passerines from the United States so that we were working with related trophically similar species across cities with relatively comparable histories. We focused on Passerines because they are a broadly comparable group with many species that have colonized cities. More precisely, we focused on native Passerines present and thought to be breeding in cities in the United States, and discarded observations of transient birds likely on route to their breeding grounds and observations of birds from outside of the breeding season. This ensured that our observations were focused on a period of high resource demand. We chose observations from May 27^th^ to July 7^th^ as our breeding season, following the practices of the North American Breeding Bird Survey (available from https://www.pwrc.usgs.gov/bbs/index.cfm). Our observations were filtered to include only those between 2010–2017 to match data availability for city variables which were last measured in 2010 and to allow for the maximum accumulation of species detections in cities. The observation dates were filtered using the R package *auk* (version 0.3.3; Strimas-Mackey et al. 2018) in R version 3.5.0 (R Core Team 2018).

We extracted species-level data for body mass, clutch size, and longevity from the Amniote life-history database (Myhrvold et al. 2016). This is a systematically compiled database of life□history traits for birds, mammals, and reptiles built for comparative life-history analyses (Myhrvold et al. 2015). Bird species that are typically present in a single U.S. state year-round were classified as residents, and birds that do not have a year-round presence in a state (i.e., migrate from elsewhere to breed) were classified as migrants. Next, we searched for data on nesting site preferences on the Cornell Lab of Ornithology Birds of North America online page (available from https://birdsoftheworld.org/bow/home) and filled in missing information with information from the Handbook of the Birds of the World (Del Hoyo et al. 2003-2011).

We assigned georeferenced eBird species records to urban areas using U.S. Census-defined urban area maps provided by the U.S. Census Bureau (U.S. Census Bureau 2010a). The urban area shapefiles define an urban area as a densely developed territory with at least 2500 people (U.S. Census Bureau 2010a). We used the R packages *sp* (version 1.3-1; Bivand et al. 2013), *rgdal* (version 1.3-4; Bivand et al. 2018), and *maps* (version 3.3.0; Becker & Wilks 2018) for this merge. Next, we calculated socioeconomic features for these urban areas using census data from the U.S. Census Bureau (U.S. Census Bureau 2010b). Data were available for the year 2010 and not the full span of our bird observation data. We thus assume that these data have remained comparable.

We were interested in species presence or absence in an urban area during a breeding season, not the number of observations of each species, and so our final data set was made up of presence data for each species observed in each city. We then excluded vagrants, introduced, and accidental observations of bird species from each state (see SI Tab. 1), as well as the brood parasites, Bronzed Cowbird (*Molothrus aeneus*) and Brown-headed Cowbird (*Molothrus ater*) from the family Icteridae, as brood parasites do not carry the fitness cost of reproduction in the same way that other species do.

### Data analyses

Human population size, housing density (housing units per square mile), and median household income were not strongly correlated (SI Fig 3) and were treated as independent variables in a series of mixed models that used species traits as dependent variables. These species-level traits were clutch size, longevity (the lifespan of an individual in years), body mass (mass of an individual in grams), migratory status (migratory or resident), and nesting habitat (tree nesting, ground nesting or cavity nesting). Data were scaled to standardize the range of independent and dependent variables to make model effects comparable.

Clutch size, longevity, and body mass were treated as dependent variables in a series of linear mixed-effects models (LMMs). All urban variables were fit in each model. We also included taxonomic family and U.S. state as random effects allowing intercepts to vary. Random intercepts estimate between-group variation in means, as well as variation within groups in each of our dependent variables. Model residuals were plotted against the expected values and we saw no strong violations of the models’ assumptions, except for body mass. We log_10_-transformed body mass to ensure the normality of residuals.

Migratory status and nesting preference were binary variables and so we fit these models using generalized linear mixed-effects models (GLMMs) with a binomial error structure and logit link function. The model structure was similar to that for LMMs with family and state treated as random effects and the city traits fit as independent variables. Migratory status was coded 1 for migratory species and 0 for residents. We used a similar series of models to explore nesting habits with tree nesters, ground nesters, and cavity nesters coded as 1 in three separate models and compared against all other nesting habits coded as 0.

## Results

Our final dataset included 44,670 observations of 160 bird species from 26 families that had been observed at least once in 1,908 cities during at least one breeding season (Fig 1). Both housing density and median income were negatively related to species longevity and body mass. There were no detectable relationships between clutch size and any socioeconomic measure and no relationship between population size and any life-history trait (Fig 2; Tab. 1). Additionally, as housing density and median income increased, the likelihood of species being migratory increased (Fig 3; Tab. 2). There was a negative relationship between housing density and cavity nesters. There were no other detectable relationships identified in our models.

**Table 1.** Model summaries and the number of observations for the urban predictors of passerine life-history traits and the predictors of passerine migratory status in the United States. One model was fit to data per response variable, including all the urban characteristics: housing density, median income, and human population size. Body mass was log10-transformed. The coefficient of variation is an indicator of model fit. Random effects were specified as (1 | family) and (1 | state), where each level of the grouping factors, family and state, had their own random intercept. The symbol σ2 is the residual variance; τ00 is the variance among the random effects; and N the total number of groups. Number of observations is the same for all the variables (n = 44,670).

| *Predictors* | **Longevity** |  | **Clutch size** |  | **Body mass** |  |
| --- | --- | --- | --- | --- | --- | --- |
|  | *Estimates* | *95% CI* | *Estimates* | *95% CI* | *Estimates* | *95% CI* |
| Intercept | -0.334 | -0.672 – 0.005 | 0.344 | -0.119 – 0.807 | 3.170 | 2.869 – 3.471 |
| Housing density | -0.034 | -0.045 – -0.024 | 0.001 | -0.007 – 0.010 | -0.011 | -0.018 – -0.005 |
| Median income | -0.022 | -0.029 – -0.014 | 0.003 | -0.002 – 0.009 | -0.011 | -0.016 – -0.007 |
| Population size | 0.001 | -0.009 – 0.010 | 0.000 | -0.008 – 0.008 | 0.004 | -0.002 – 0.010 |
| **Random Effects** |  |  |  |  |  |  |
| $\sigma^2$ | 0.50 | | 0.31 | | 0.20 | |
| $\tau_{00}$ | 0.01<sub>State</sub> | | 0.00<sub>State</sub> | | 0.00<sub>State</sub> | |
|  | 0.75<sub>family</sub> |  | 1.44<sub>family</sub> |  | 0.60<sub>family</sub> |  |
| N | 26<sub>family</sub> |  | 26<sub>family</sub> |  | 26<sub>family</sub> |  |
|  | 49<sub>State</sub> |  | 49<sub>State</sub> |  | 49<sub>State</sub> |  |

**Table 2.** Model summaries for urban predictors of passerine migratory status and nesting preferences in the United States. One model was fit to data per response variable, including all the urban characteristics. The coefficient of variation is an indicator of model fit. Species are classed as migratory = 1 or resident = 0; tree nesting, ground nesting and cavity nesting = 1 or other places = 0. An odds ratio greater than one indicates that the chance of finding a migrant or a bird with a specific nesting preference is higher than the chance of finding a resident or a species with any other nesting preference. Random effects were specified as in Table 1. The symbol σ2 is the residual variance; τ00 is the variance among the random effects; and N the total number of groups. Number of observations is the same for all the variables (n = 44,670).

|  | **Migratory status** |  | **Tree nesting** |  | **Ground nesting** |  | **Cavity nesting** |  |
| --- | --- | --- | --- | --- | --- | --- | --- | --- |
| *Predictors* | *Odds Ratios* | *95 % CI* | *Odds Ratios* | *95% CI* | *Odds Ratios* | *95% CI* | *Odds Ratios* | *95% CI* |
| Intercept | 1.551 | 0.053 – 45.176 | 0.000 | 0.000 – 0.018 | 0.046 | 0.002 – 1.317 | 0.000 | 0.000 – 0.000 |
| Housing density | 1.141 | 1.087 – 1.198 | 1.035 | 0.993 – 1.079 | 1.002 | 0.959 – 1.046 | 0.918 | 0.864 – 0.975 |
| Median income | 1.098 | 1.064 – 1.133 | 0.997 | 0.968 – 1.026 | 1.015 | 0.983 – 1.048 | 0.973 | 0.935 – 1.014 |
| Population size | 1.026 | 0.975 – 1.079 | 0.977 | 0.940 – 1.015 | 1.011 | 0.970 – 1.054 | 1.015 | 0.957 – 1.075 |
| **Random Effects** |  |  |  |  |  |  |  |  |
| $\sigma^2$ | 0.65 | | 0.68 | | 0.57 | | 0.40 | |
| $\tau_{00}$ | 0.21<sub>State</sub> | | 0.01<sub>State</sub> | | 0.01<sub>State</sub> | | 0.06<sub>State</sub> | |
|  | 92.60<sub>family</sub> |  | 161.40<sub>family</sub> |  | 84.77<sub>family</sub> |  | 508.98<sub>family</sub> |  |
| N | 26<sub>family</sub> |  | 26<sub>family</sub> |  | 26<sub>family</sub> |  | 26<sub>family</sub> |  |
|  | 49<sub>State</sub> |  | 49<sub>State</sub> |  | 49<sub>State</sub> |  | 49<sub>State</sub> |  |

**Figure 1.**
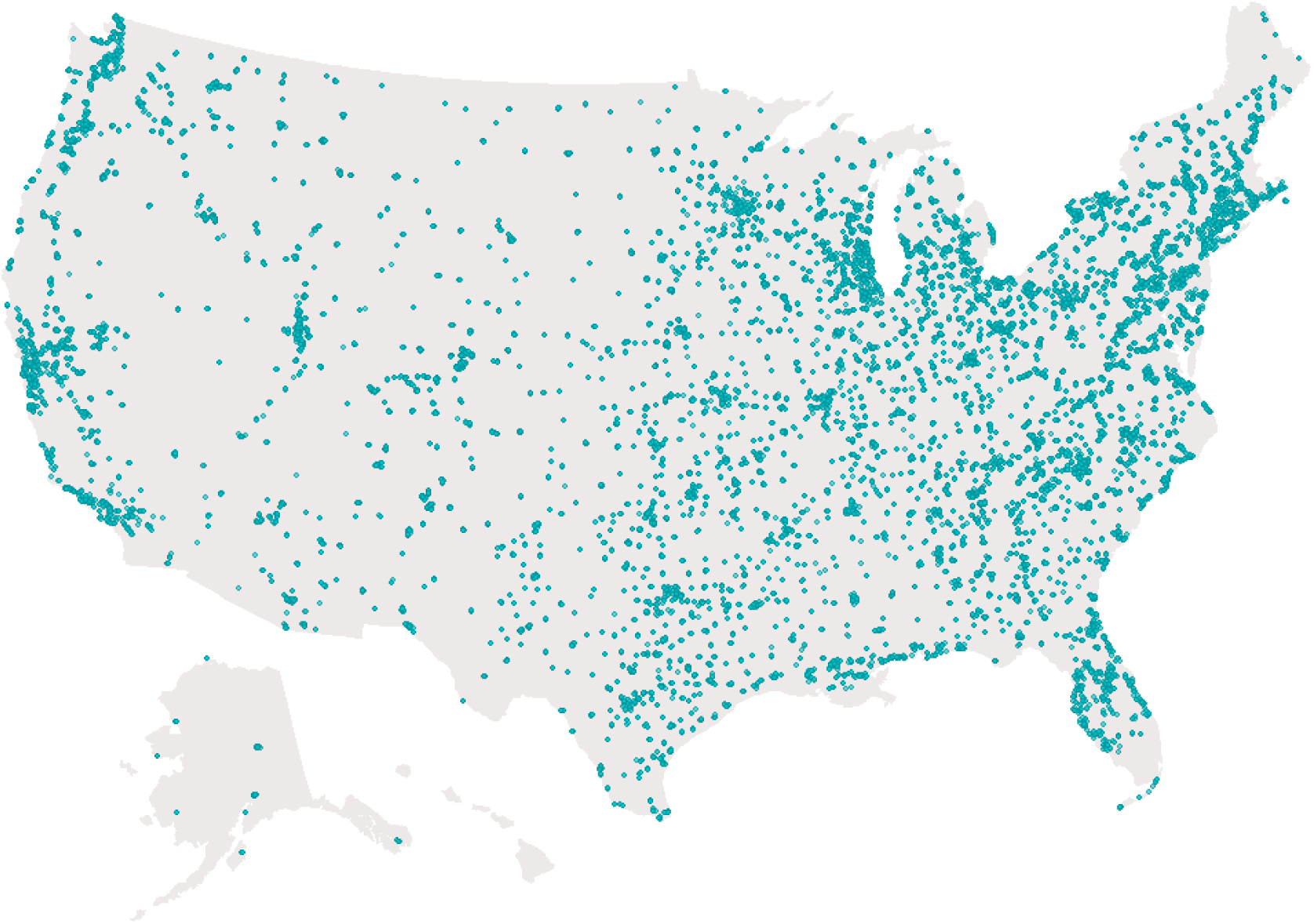
Map of 44,670 bird observations of 160 Passerine species in 1,908 urban areas across the United States. Circles represent the longitude and latitude coordinates of bird observations.

**Figure 2.**
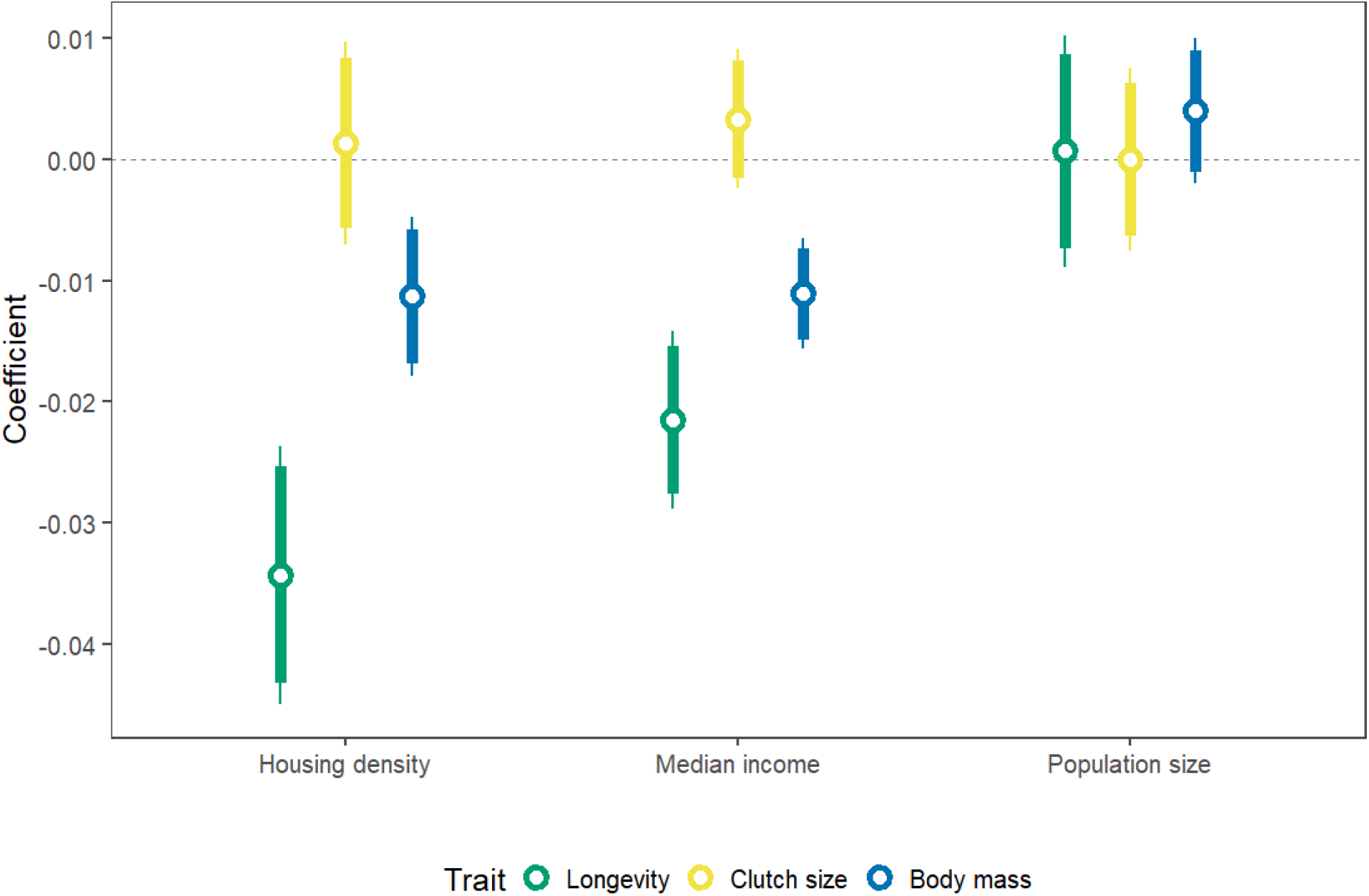
Linear mixed-effects model coefficients for urban predictors of passerine life-history traits in the United States. Open circles represent coefficient estimates, bold whiskers are 90% confidence intervals, and narrow whiskers are 95% confidence intervals. Body mass was log_10_ - transformed. Sample size is the same for all variables (n = 44,670).

**Figure 3.**
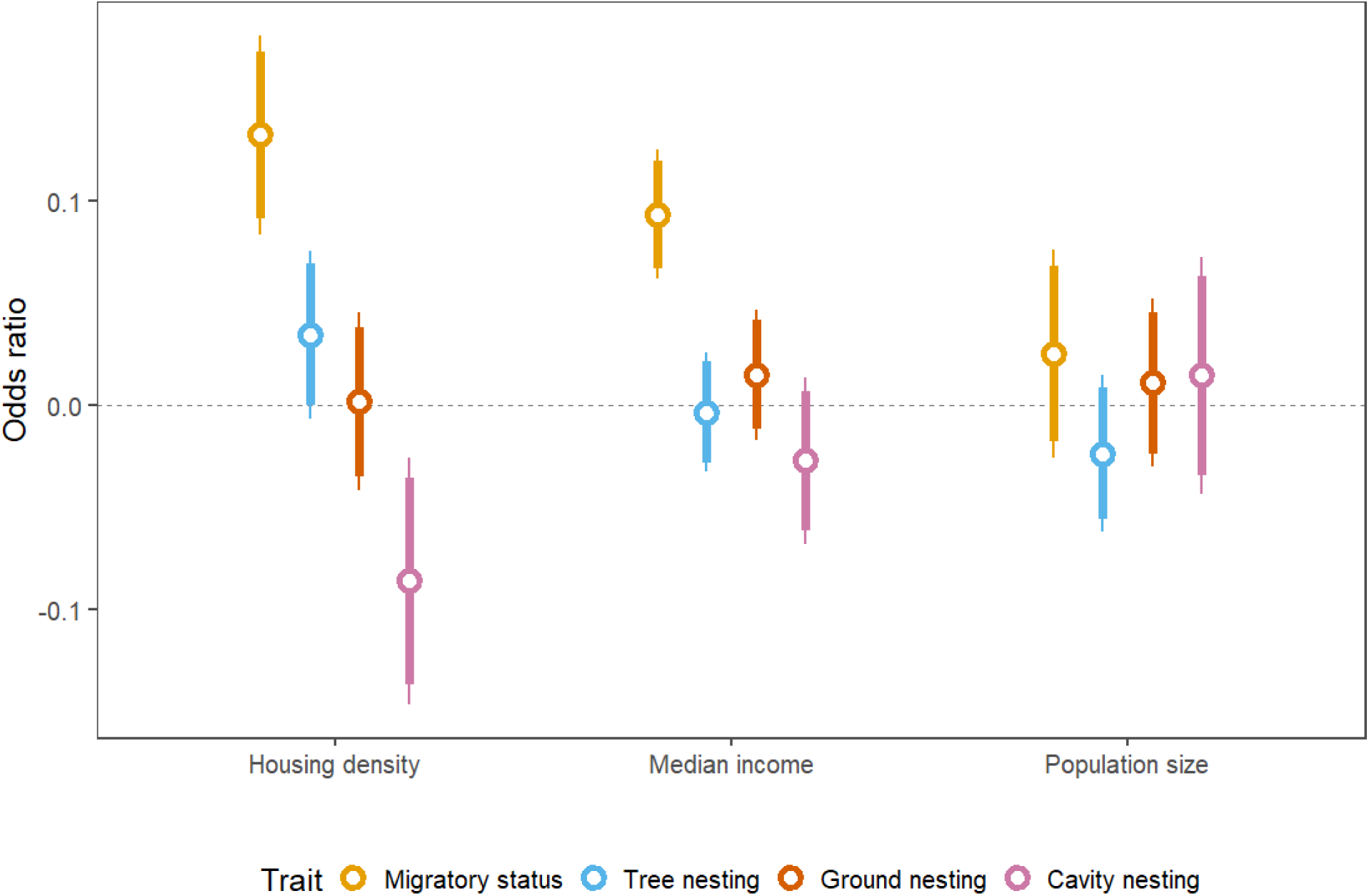
Generalized linear mixed-effects model coefficients for the predictors of passerine migratory status and nesting preferences in the United States. Open circles are coefficient estimates, bold whiskers are 90% confidence intervals, and narrow whiskers are 95% confidence intervals. Species are classed as migratory = 1 or resident = 0; tree nesting, ground nesting and cavity nesting = 1 or other places = 0. An odds ratio greater than one indicates that the chance of finding a migrant bird or a species with a certain nesting preference in the area is higher than a chance of finding a resident or a bird with any other nesting preference, and vice versa. The sample size is the same for all the variables (n = 44,670).

## Discussion

The typical lifespans, body masses, and migratory status of species found across different cities varied with housing density and median income (Fig 4). This presumably reflects differences in the average fitness of a species given the local features of a particular city. This suggests that ecological selection—in this case associated with aspects of socioeconomics—plays an important role in determining biodiversity at the earliest stage of community formation following the emergence of new environments. Like environmental variation in natural habitats, cities vary in the details of their composition and different life-history strategies are better suited to different types of city.

**Figure 4.**
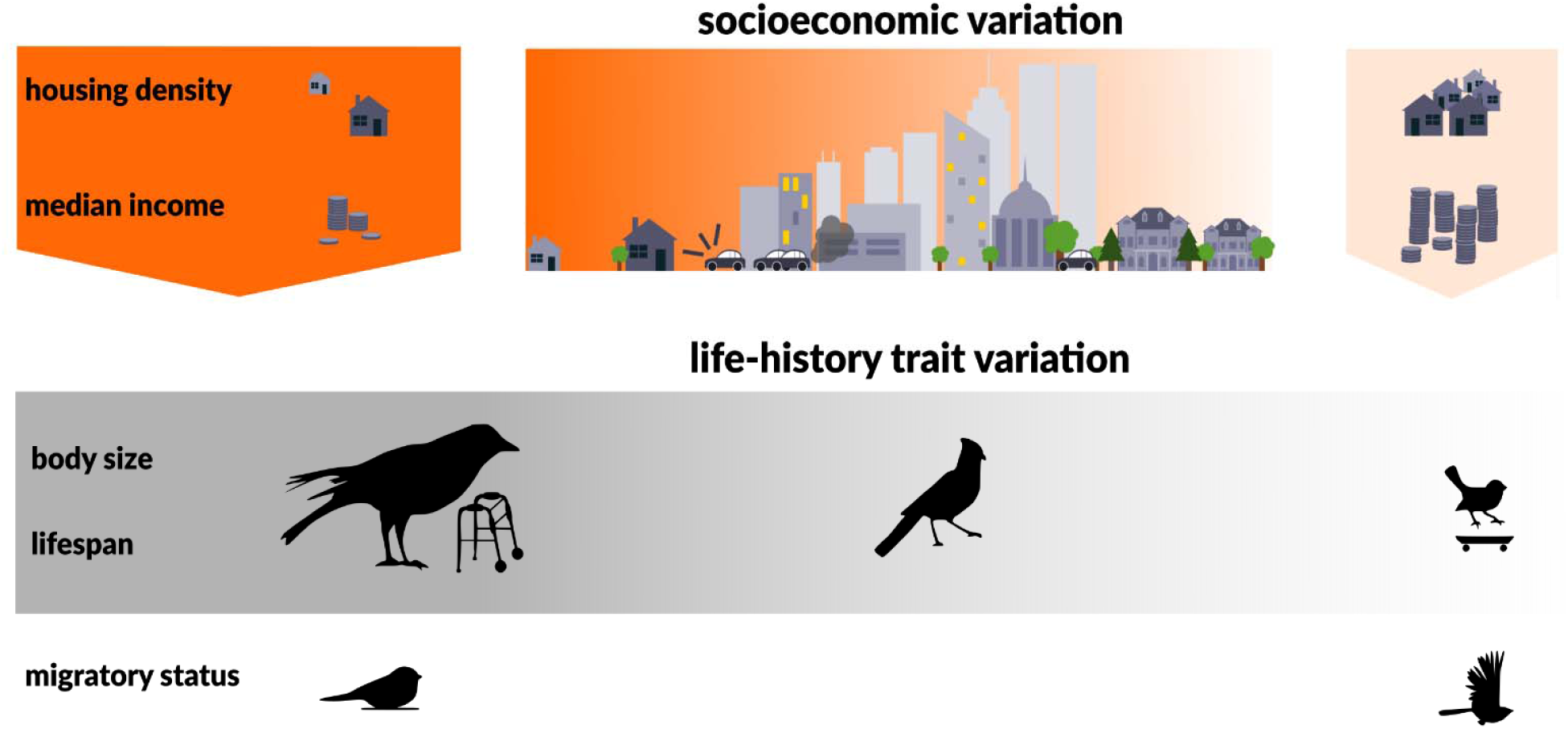
Our results suggest that from a bird’s eye view cities are heterogeneous habitats supporting different life-history strategies that depend on the socioeconomic variation across cities. As a city’s housing density and median income increased it supported species with smaller body sizes and shorter lifespans, and more migratory species. This suggests that differential survival and reproduction at the species-level can quickly generate geographical patterns of species trait variation across urban environments similar to those found in natural settings without the need for evolutionary change.

It is generally expected that as the typical body size of a species increases, its reproductive output decreases, and lifespan increases (Bielby et al. 2007). This trait combination is thought to reflect trade-offs between current and future reproduction and survival given resource availability. We found that species body mass and lifespan varied together as expected if cities with high housing density and median income were relatively stable resource-rich environments (see description below). As the median income and housing density of a city increased the city tended to support smaller shorter-lived species. Contrary to our predictions, we found no evidence that clutch size covaried with either these life-history traits or the socioeconomic status of a city. Recent explorations of life-history trait variation have also found that mean number of offspring was not well correlated with other traits associated with the fast–slow continuum for birds and suggested that annual reproductive effort represents a second life-history strategy axis of variation (Healy et al. 2019). This could happen if trade-offs associated with reproduction do not always result in changes in annual reproductive output (Healy et al. 2019). More natural areas with plentiful and stable resources tend to support greater numbers of small-bodied species that prioritize reproduction at the expense of lifespan, due to high offspring survival rates. Stochastic or resource-poor environments tend to support larger, longer-lived species that prioritize adult survival over producing many offspring. This strategy is thought to buffer against the consequences of reproductive failures by spreading the risk of offspring mortalities across multiple breeding attempts during a longer lifetime. If this interpretation holds for different cities, it suggests that cities with high housing density and median income may be relatively stable, predictable habitats with readily available resources for urban birds. Our results further suggest that human population size is not a strong ecological filter for these traits.

The results for migratory species support our resource and environmental variability-based interpretation of a city’s housing density and median income. Urban areas with higher median income and greater housing densities tended to support relatively more migratory species, whereas again the human population size of a city was not important. Migratory species breed in cities less often than natural areas (Poague et al. 2000; Croci et al. 2008) and generally choose breeding sites with an excess of resources that are not completely exploited by resident birds (Dalby et al. 2014; Somveille et al. 2015). That more migratory species choose to breed in cities with high housing densities and higher median incomes suggests that they may be choosing these cities, at least in part, based on resource availability (Martin & Karr 1986; Faaborg et al. 2010; Jenkins et al. 2017).

Cities with more economic resources and higher housing densities might provide more stable, high-quality habitats for birds in many ways. There is a well-recognized positive correlation between urban plant diversity, vegetation cover, and wealth in cities (Iverson & Cook 2000; Hope et al. 2003; Kinzig et al. 2005; Leong et al. 2018). Higher housing density typically also means more backyards and gardens in people’s yards in a city, and these diverse plant communities can provide important sources of food, cover, and nesting resources for birds (Thompson et al. 2003; Smith et al. 2005, 2006; Narango et al. 2018). In addition to vegetation, cities also provide stable year-round access to food sources via bird feeders and human food waste (Lepczyk et al. 2004; Tryjanowski et al. 2015) which influences urban bird community structure and breeding success (Robb et al. 2008; Galbraith et al. 2015). Wildlife feeding is a popular activity among people with recent estimates of ∼ 59% of U.S. households providing food for mainly birds (U.S. Fish & Wildlife Service 2016). Bird feeders are also more concentrated in densely populated areas of higher socioeconomic class (Fuller et al. 2008, 2013). Bird feeders can act as particularly important resources to Passerines, as many species are mainly granivores. Corvids, on the other hand, can as omnivores easily exploit human food waste. Our results also showed that as the housing density of a city increased, it tended to support fewer cavity nesting species. This suggests that fewer cavity-bearing trees and snags may be present in these cities, and that—for most species—artificial cavities and other human-associated structures cannot replace more natural nesting resources (Blewett & Marzluff 2005).

Interestingly, housing density and human population size were only moderately correlated in our data, yielding more nuanced insights into the effects of human activity versus city structure. Traits fell along our housing density axis similarly to median income, indicating that housing density, like wealth, creates stability in resources and habitat. We speculate that highly populated cities are the most disturbed in terms of traffic levels, noise, artificial light, and pollution (McKinney 2001, 2002; Luck 2007; Isaksson 2018; Strohbach et al. 2019). Each of these factors could make highly populated areas generally poor habitat for birds.

Explaining the general geographical patterns of species trait variation (e.g., body size variation) has been difficult, and the topic remains controversial (Blackburn et al. 1999; Meiri & Dayan 2003; Meiri & Thomas 2007; Olson et al. 2009). For example, body size in birds has been linked to temperature, resource availability, seasonality, and environmental stability (James 1970; Murphy 1985; Olson et al. 2009; Sun et al. 2017). Our results demonstrate that ecological selection can quickly generate patterns of species trait variation across cities similar to those found in natural settings. In this case, we did not find gradual spatial clines, but rather heterogeneous patches across which life-history traits vary similarly to those seen across natural environments. Although cities are generally more similar to each other than to their natural surroundings, our results suggest that from a bird’s eye view, they are heterogeneous habitats supporting different life-history strategies that in part depend on socioeconomic factors and the built environment. When using cities as replicates in urban in studies of urban wildlife (Szulkin et al. 2020) the assumption that different cities are directly comparable should be made with care. As the world’s most rapidly growing ecosystem, understanding what kinds of species initially colonize cities provides important information about how the distribution of biodiversity will change following rapid, human-caused environmental shifts in an era of global change. It is also important for us to understand how cities support nature because it is now within cities that most people interact with and benefit from local biodiversity (e.g., through recreational activities and ecosystem services; Bolund and Hunhammar 1999). Finally, although cities tend to support fewer species than nearby natural areas (Chace & Walsh 2006), on average urban biodiversity is primarily comprised of native species (Aronson et al. 2014). These findings in addition to our results suggest that cities could play a more important role in conservation and management than they currently do.

## Acknowledgments

We particularly want to thank Alyssa Garrard for collecting data on Passerine nesting site preferences as well as the other members of the Population Ecology & Evolutionary Genetics Group for their helpful comments on manuscript drafts. We also want to acknowledge the work done by the eBird citizen-science observers. This study was supported by a discovery grant of the Natural Sciences and Engineering Research Council of Canada (NSERC) to CJG. RPK and CS were additionally supported by the University of Manitoba Graduate Fellowships and a University of Manitoba Graduate Enhancement of Tri-council funding grant to CJG.

## Notes

### Competing Interest Statement

The authors have declared no competing interest.

### Summary of Updates

Corrected author name spelling.

